# Production of the sesquiterpene bisabolene from one- and two-carbon compounds in engineered *Methanosarcina acetivorans*

**DOI:** 10.1101/2024.09.23.614462

**Authors:** Andrea Mentrup, Luca V. Scheitz, Theo Wallenfang, Michael Rother

**Affiliations:** Fakultät Biologie, Technische Universität Dresden, Dresden, Germany

**Author notes:** Corresponding Author: Michael Rother, Fakultät Biologie, Technische Universität Dresden, 01062 Dresden, Germany. Max Delbrück Center for Molecular Medicine, Berlin, Germany (Theo Wallenfang).

**Keywords:** *Methanosarcina*, isoprenoids, bisabolene, mevalonate pathway, engineering

## Abstract

The isoprenoid bisabolene, one of the simplest monocyclic sesquiterpenes, is a natural plant product that, in addition to its biological function, serves as a precursor for many industrial products. Due to the low concentration of bisabolene and the long harvest cycle, industrial production of this isoprenoid in plants is economically challenging. Chemical synthesis of bisabolene also suffers from significant disadvantages, such as low yields, toxic side products and high costs. Archaea appear suitable producers of isoprenoids, as their membrane lipids consist of isoprenoid ethers, which are synthesized via a variant of the mevalonate pathway. Archaeal model species have versatile metabolic capacities, which makes them potential candidates for biotechnological applications. Here, we engineered *Methanosarcina acetivorans* for production of α-bisabolene from one-carbon substrates by introducing a bisabolene synthase from *Abies grandis*. Expression of a codon-optimized bisabolene synthase gene in *M. acetivorans* resulted in 10.6 mg bisabolene/L of culture. Overexpressing genes of the mevalonate pathway only slightly increased bisabolene yields, which, however, were reached much earlier during incubations than in the corresponding parent strain. The data presented argue for the suitability of *M. acetivorans* for the biotechnical production of certain isoprenoids.

## Introduction

Isoprenoids are natural products composed of C5-isoprene units and are among the most diverse compounds synthesized by biological systems. With more than 35,000 known isoprenoids, these compounds vary in size, structure, and biological function. They play essential roles in various cellular processes, including regulation of gene expression, energy conservation, cell membrane structure, signaling, cell defense, and as vitamins (Chandran et al., 2011). Beyond their biological functions, isoprenoids have numerous biotechnological applications, as biofuels, insecticides, pharmaceuticals, colorants, flavorings, fragrances, and precursors for semisynthetic approaches (Pfeifer et al., 2021; Lei et al., 2022). Depending on the type of substance required, there are various strategies for meeting the increasing demand for economically relevant isoprenoids. The first option is their extraction from plant material, which, however, faces various challenges. On the one hand, the isolation of isoprenoids produced by plants often requires substantial energy and resource consumption. On the other hand, the quality of the product fluctuates due to different environmental conditions (Chang and Keasling 2006; Immethun et al., 2013). Alternatively, isoprenoids can be produced by chemical synthesis from petrochemical raw materials. Beside the problem of finiteness of the resource, the complexity and thus the yield pose a problem. It also requires significant amounts of energy while producing toxic by-products (Chemler et al., 2006; Winter and Tang 2012). Thus, biotechnical approaches represent favorable tracks for the production of isoprenoids. Microorganisms exhibit rapid growth and can utilize inexpensive carbon sources to produce products without the generation of toxic by-products, thereby offering a cost-effective and environmentally benign production process (Chemler et al., 2006; Ajikumar et al., 2008).

Bisabolene is one of the simplest monocyclic sesquiterpenes (Souček et al., 1961) and is characterized by a remarkable diversity of applications in various industrial branches. Despite its low concentration in plants, which makes economically viable isolation from plant sources difficult, bisabolene has considerable industrial potential (Zhao et al., 2021). This low availability in nature means that alternative production methods, such as biotechnological approaches, are becoming increasingly important. In the cosmetics and perfume industry, bisabolene is employed as a fragrance and flavoring agent due to its pleasant and balsamic-woody aroma (Wang et al., 2017). Another, particularly innovative, application of bisabolene is as a starting material for the biosynthesis of bisabolane, a potential alternative fuel to D2 diesel. This application illustrates the potential of bisabolene to contribute to the development of sustainable and renewable energy sources (Peralta-Yahya et al., 2011).

The mevalonate (MVA) and methylerythritol phosphate (MEP) pathways represent two distinct biochemical pathways for the synthesis of isoprenoids, which are present in a vast array of organisms. The MEP pathway is utilized by a multitude of bacteria, plants, and algae for the synthesis of the isoprenoid precursor isopentenyl diphosphate (IPP) and its isomer dimethylallyl diphosphate (DMAPP) (Zhao et al., 2013). In contrast, the MVA pathway represents the conventional route for the synthesis of isoprenoids in archaea, eukaryotes, and some bacteria. Studies on *Methanocaldococcus jannaschii* and *Aeropyrum pernix* established a modified version of the canonical eukaryotic MVA pathway, which is present in most members of the phylum Archaea and is called the archaeal MVA pathway (Fig. 1) (Grochowski et al., 2006; Hayakawa et al., 2018). The archaeal MVA pathway, which is also present in *Methanosarcina* species, requires the enzyme acetoacetyl-CoA thiolase, which catalyzes the condensation of two acetyl-CoA molecules to form acetoacetyl-CoA. Subsequently, HMG-CoA synthase combines two acetoacetyl-CoA molecules to produce 3-hydroxy-3-methylglutaryl-CoA (HMG-CoA), which is then reduced to mevalonate by HMG-CoA reductase (Miziorko 2011). Mevalonate is phosphorylated to mevalonate-5-phosphate by mevalonate kinase. Subsequently, the phosphomevalonate dehydratase catalyzes the dehydration of mevalonate-5-phosphate to trans-anhydromevalonate 5-phosphate, which is then decarboxylated to isopentenyl phosphate by anhydromevalonate phosphate decarboxylase (AMPD). Prenylated flavin mononucleotide synthase plays a pivotal role in the synthesis of prenylated flavin mononucleotide, which is postulated to be the essential cofactor for the activity of AMPD. Isopentenyl phosphate is phosphorylated to IPP by isopentenyl phosphate kinase. IPP, the principal isoprenoid building block, is isomerized to DMAPP. Both IPP and DMAPP serve as precursors for synthesizing more complex isoprenoids (Yoshida et al., 2020).

**Figure 1:**
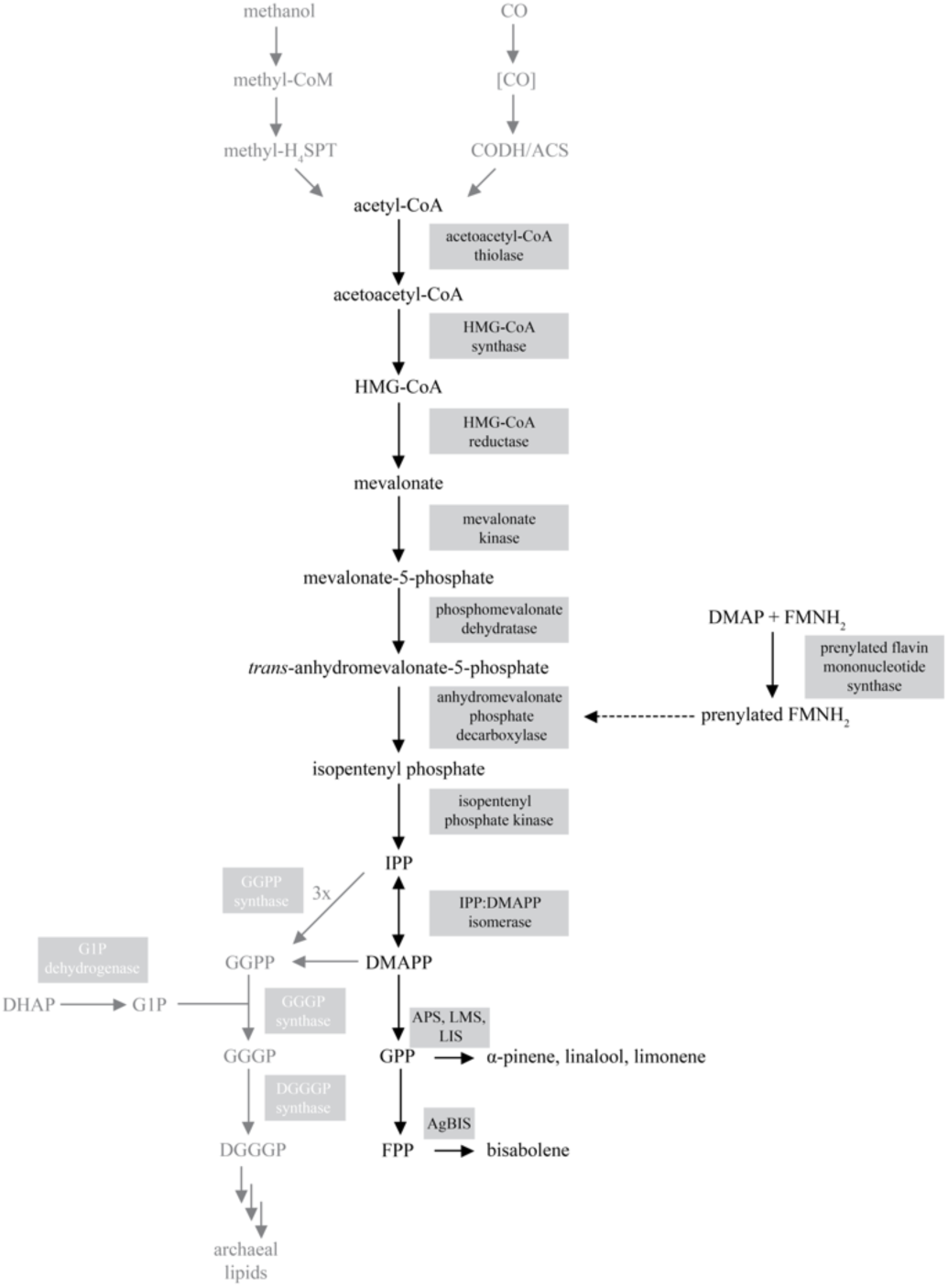
Scheme of the archaeal mevalonate pathway. Archaeal membrane are composed of isoprene subunits linked to glycerol-1-phosphate via ether bonds. The isoprene precursors IPP and DMAPP are synthesized from acetyl-CoA (generated for anabolic purposes from methanol and during catabolism from CO) via the mevalonate (MVA) pathway. The enzymes involved are boxed; abbreviations: CoA, coenzyme A; HMG, 3-hydroxy-3-methyl-glutaryl; DMAP, dimethylallyl phosphate; FMNH_2_, reduced flavin mononucleotide; IPP, isopentenyl pyrophosphate; DMAPP, dimethylallyl pyrophosphate; GPP, geranyl pyrophosphate; GGPP, geranylgeranyl diphosphate; FPP, farnesyl pyrophosphate; DHAP, dihydroxyacetone phosphate; G1P, *sn*-glycerol-1-phosphate; GGGP, 3-*O*-geranylgeranyl-*sn*-glyceryl-1-phosphate; DGGGP, 2,3-bis-*O*-geranylgeranyl-*sn*-glyceryl-1-phosphate; IPS, isoprenoid synthase.

*Methanosarcina acetivorans* belongs to the methanogenic archaea, the only organisms that produce methane (CH_4_) as means of their energy metabolism. These archaea are essential for the complete remineralization of organic material in oxygen-depleted environments (Whitman et al., 2006). Species of the genus *Methanosarcina* are characterized by their metabolic versatility. They can utilize a variety of one- and two-carbon substrates for growth, including methanol, methylamines, methyl sulfides, acetate and carbon monoxide (CO), to produce methane (Schöne and Rother 2018).The latter capacity might be a particular advantage that makes *M. acetivorans* a good candidate for isoprenoid production because large amounts of acetate are formed during growth on CO (Rother and Metcalf 2004). Since the isoprenoid precursors IPP and DMAPP and acetate are generated from acetyl-CoA, catabolic levels of the starting material should be available in the cell. Furthermore, archaea do not employ the MVA pathway for the formation of secondary metabolites, but for the synthesis of their membrane lipid precursors (Jain et al., 2014), requiring a comparably high turnover. For this purpose, the isoprenoid precursors IPP and DMAPP are converted to geranylgeranyl diphosphate. Together with glycerol-1-phosphate, which originates from glycolysis/gluconeogenesis, the enzyme geranylgeranyl-glyceryl-1-phosphate (GGGP) synthase catalyzes the synthesis of GGGP. This product is incorporated into archaeal lipids through further steps (Fig. 1). Thus, the prospect of *M. acetivorans* being able to produce IPP not only in trace amounts makes it a promising candidate for isoprenoid production. Indeed, this organism produced up to 850 mg/L of isoprene when the gene for isoprene synthase was heterologously expressed (Aldridge et al., 2021).

Although the biochemical and bioenergetic aspects of methanogenic energy metabolism are well understood and sophisticated tools for their genetic manipulation are available for some model methanogens, their industrial application is currently limited to only a few areas. Their ability to produce methane is used either in biogas plants, anaerobic wastewater treatment or in power-to-gas concepts (Angenent et al., 2018; Enzmann et al., 2018; Contreras et al., 2022). Moreover, archaea have been employed in the food production industry as protein producers. Furthermore, individual archaeal enzymes are already being employed in industrial processes due to their adaptation to extremophilic conditions. The industrial enzymes include proteases, lipases, amylases, cellulases as well as polymerases, DNA and RNA ligases and endonucleases. In addition to these already established applications, various efforts are being made to further utilize archaea for biotechnological processes. Some examples of this are fuel production, as well as the production of hydrogen, compatible solutes, lipids, bacteriorhodopsin and carotenoids (Pfeifer et al., 2021; Aparici-Carratalá et al., 2023), To further exploit potential applications for methanogenic archaea, in this study we have monitored production of various isoprenoids by genetically modified *M. acetivorans* strains. A special focus was on α-bisabolene, for which the use of different energy substrates and different cultivation regimes was studied.

## Materials and Methods

### Strains, growth, and transformation

*Escherichia coli* strains DH10B (Grant et al., 1990) and WM1788 (for propagating R6K replicons) (Haldimann and Wanner 2001) were grown and transformed employing standard conditions (Ausubel et al., 2003). *Methanosarcina acetivorans* strains, listed in Table S1, were cultivated in high salt (HS) medium (Metcalf et al., 1996) containing trace elements and vitamins (Sowers and Noll 1995). Either 26 mL glass (Balch) tubes (Ochs, Bovenden, Germany) containing 10 mL or 5 mL of medium, or 100 mL serum bottles (Ochs) containing 50 mL of medium were used. 125 mmol/L methanol (Sigma-Aldrich, Munich, Germany) plus 40 mmol/L sodium acetate (Carl Roth, Karlsruhe, Germany), supplemented from anaerobic, sterile stock, were used as growth substrate, or 150 kPa carbon monoxide (CO) (Air Liquide, Düsseldorf, Germany), added through sterile filters. For CO-dependent cultivation, the medium contained 60 mM piperazine-N,N′-bis(2-ethanesulfonic acid) (PIPES), pH 7.4, as buffer (Schöne et al., 2022). For induction of the tetracycline-inducible promoters, the media contained 100 µmol/L tetracycline (tet) (Carl Roth). Growth was monitored photometrically at 578 nm (OD_578_) either directly (i.e., undiluted) in Balch tubes using a Genesys 20 spectrophotometer (Thermo Scientific, Langenselbold, Germany) or using an Ultrospec 2000 spectrophotometer (Pharmacia Biotech, Uppsala, Sweden) for measurements of diluted culture samples. In the presence of different isoprenoids growth of *M. acetivorans* was monitored nephelometrically using a cell growth quantifier (CGQ) (Aquila Biolabs GmbH, Baesweiler, Germany) (Bruder et al., 2016) in serum bottles with 50 mL media as described (Richter et al., 2024). The bisabolene-producing *M. acetivorans* strains were transferred for two or three times to medium supplemented with tet and, when quantification of the isoprenoid was intended, overlaid with dodecane (10% v/v, Sigma-Aldrich). Since each culture had to be sacrificed for sample extraction (see below), two to five independent cultures represent each sample point.

Strains of *Methanosarcina acetivorans* (listed in Table S1) were transformed as described (Oelgeschläger and Rother 2009). Markerless deletion of *ssu*, MA2965, or MA3852, concomitant with inserting *trans*-genes was conducted as described for *hpt* (Pritchett et al., 2004).

### Generation of plasmids

Plasmids used in this study are listed in Table S2. DNA fragments deriving from PCR (oligonucleotides used are listed in Table S3) and used for cloning were sequenced by Microsynth Seqlab using the BigDye Terminator Cycle Sequencing protocol.

Through a review of the literature on isoprenoid synthases from plants and bacteria, we chose the monoterpenes α-pinene, limonene, and linalool to be produced via α-pinene synthase from *Pinus taeda*, limonene synthase from *Mentha spicata*, and linalool synthase from *Streptomyces clavuligerus* for production in *M. acetivorans* (Table S4). The genes were retrieved from GenBank (NCBI accession numbers are given in Table S4), trimmed of plastid localization signals, and codon-optimized for *M. acetivorans* using Geneious Prime (v. 2019.2.3 Biomatters Ltd., Auckland, New Zealand) using a custom codon usage table; the sequences were subsequently manually adjusted to a codon adaptation index of 0.6-0.8 (Sharp and Li 1987). The fusion of the tet-inducible promoter PmcrB(tetO1) (Guss et al., 2008) and the respective isoprenoid synthase gene was obtained commercially. The synthesis products were then transferred to the *M. acetivorans/E. coli* shuttle vector pMR08 (Oelgeschläger and Rother 2009) via SphI and SpeI. In the case of AgBIS (Table S2), the synthesized plasmid was first used as a template for PCR amplification with the primers o_SphI_pmcrB_tetO1_for and o_SpeI_HindIII_AgBIS_rev (Table S3) to remove the His_6_-tag encoding sequence, before moving the fusion to pMR08.

The ten genes encoding mevalonate pathway enzymes from *M. mazei* were divided into three modules, in order to facilitate cloning and to identify potential bottlenecks of carbon flow through the pathway (Table S4). Module MVA I consists of the genes MM_0870, MM_0871 and MM_0335. MVA II consists of the genes MM_1526, MM_1525, MM_1524 and MM_1871. MVA III consists of the genes MM_1762, MM_1763, MM_1764 (Table S4). In the case of MVA I and MVA II, all genes were combined into a synthetic operon and the fusion product was synthesized fused to a tet-inducible promoter. The synthesis product was then moved to pMssu in the case of MVA I, and to the plasmid pMA_3852 in the case of MVA II via NotI. Since the genes assembled in the MVA III module are present in a natural operon of *M. mazei*, the entire operon was amplified by PCR from the *M. mazei*\* genome (Ehlers et al., 2005) using the primers o_NdeI_Operon_MM1762_1764_for and o_Operon_MM1762_1764_BamHI_rev (Table S3). The resulting PCR product was cloned into plasmid pJK028a via NdeI and BamHI, thereby creating a PmcrB(tetO3) fusion. The fusion was PCR amplified using the primers o_NotI_pmcrB_tetO3_Operon_MM1762_64_for and o_pmcrB_tetO3_Operon_MM1762_64_NotI_rev, and the resulting PCR product was cloned into plasmid pMA2965 via NotI.

### Analysis of bisabolene

For the determination of bisabolene concentrations from culture supernatants, individual cultures were sacrificed, as the dodecane phase of the respective culture was removed and centrifuged at 10,000 g for 5 min. 200 µL of the organic phase was transferred to an air-tight vial and 2 µL was analyzed by gas chromatography as described (Zhao et al., 2021). Briefly, a GC-2010 Plus gas chromatograph (Shimadzu Europa GmbH, Duisburg, Germany) was operated with nitrogen as the carrier gas at an inlet pressure of 124.6 kPa, and equipped with an Optima 5 column (Macherey Nagel, Dueren, Germany). The temperature of the GC oven was initially maintained at 60 °C for 3 min and then increased to 280 °C at a rate of 12 °C per min. This temperature was then held for 5 min. The injector was held at 300 °C and the temperature of the flame ionization detector was also set at 300°C. Data were recorded and analyzed using lab solution software version 5.98 (Shimadzu Europa GmbH, Duisburg, Germany). A bisabolene isomer mixture (Thermo Fisher scientific, Darmstadt, Germany) was used to generate a calibration curve by adding known amounts to culture tubes, overlaying with dodecane, and treating the standards in the identical fashion as the analytes (described above). For statistical analysis, GraphPad Prism software (version 5.03) was employed. The statistical tests employed are highlighted in the respective figure legends.

## Results and Discussion

### Production of monoterpenes in M. acetivorans

In a previous study, *M. acetivorans* was genetically amended by introducing the gene for isoprene synthase from *Populus alba*, which resulted in a strain producing the hemiterpene isoprene (Aldridge et al., 2021). We wanted to extend on this finding and genetically engineer *M. acetivorans* to synthesize larger isoprenoids. To achieve this, we introduced genes for α-pinene synthase from *Pinus taeda*, limonene synthase from *Mentha spicata*, and linalool synthase from *Streptomyces clavuligerus*, respectively, into *M. acetivorans* (Table S4). The codon usage of the *trans*-genes was adapted to that of *M. acetivorans*, and the corresponding gene was fused with to a well-established strong promoter established inducible by tet (Guss et al., 2008). The gene fusion was placed onto an autonomously replicating vector, and the resulting plasmid was subsequently transferred into *M. acetivorans* WWM73, the parent strain used in this study. Transformants obtained were analyzed for the presence of the respective isoprenoids. However, none of the isoprenoids in question could be detected, regardless whether gene expression had been induced by tet, or not. Extraction of the isoprenoids from the medium by growing the cultures in the presence of an overlay of paraffin oil or dodecane (10%, v/v), which should also have a concentrating effect, was also attempted. However, in none of the cultures could the respective isoprenoid be detected.

The exact reason why none of the monoterpenes was produced was not thoroughly investigated and, thus, remains unclear. Still, considering the antimicrobial property of isoprenoids (Wang et al., 2019), together with a certain “leakiness” of the inducible promoter employed (Guss et al., 2008), it is possible that the producing phenotype was selected against and quickly lost. To gather some general evidence for this notion, the antimicrobial activity of several isoprenoids on *M. acetivorans* was investigated. To this end, α-pinene, linalool, geraniol, terpineol, borneol, and bisabolene, were added to the medium at varying concentrations (0.1 mmol/L, 0.5 mmol/L, 1 mmol/L, and 2.5 mmol/L) and growth of *M. acetivorans* on methanol plus acetate was assessed by means of CGQ measurements (see Materials and Methods, Fig. 2). α-pinene completely inhibited growth of *M. acetivorans* already at 0.5 mmol/L (Fig. 2A), indicating a profound antimicrobial effect on the organism. A similar effect was observed using 0.5 mmol/L geraniol, with only minimal cell proliferation detectable (Fig. 2B). Linalool (Fig. 2C), terpineol (Fig. 2D), and borneol (Fig. 2F) required 1 mmol/L to be present for the same effect. Notably, bisabolene exhibited the least severe antimicrobial effect on *M. acetivorans*. In the presence of 2.5 mmol/L, *M. acetivorans* still grew, albeit at a slower rate than without the isoprenoid (Fig. 2E). Thus, all six isoprenoids tested had a negative impact on the growth of *M. acetivorans*, with bisabolene exerting the least pronounced. Whether this effect is (partly) due to the solubility of the respective isoprenoid, was not investigated here.

**Figure 2:**
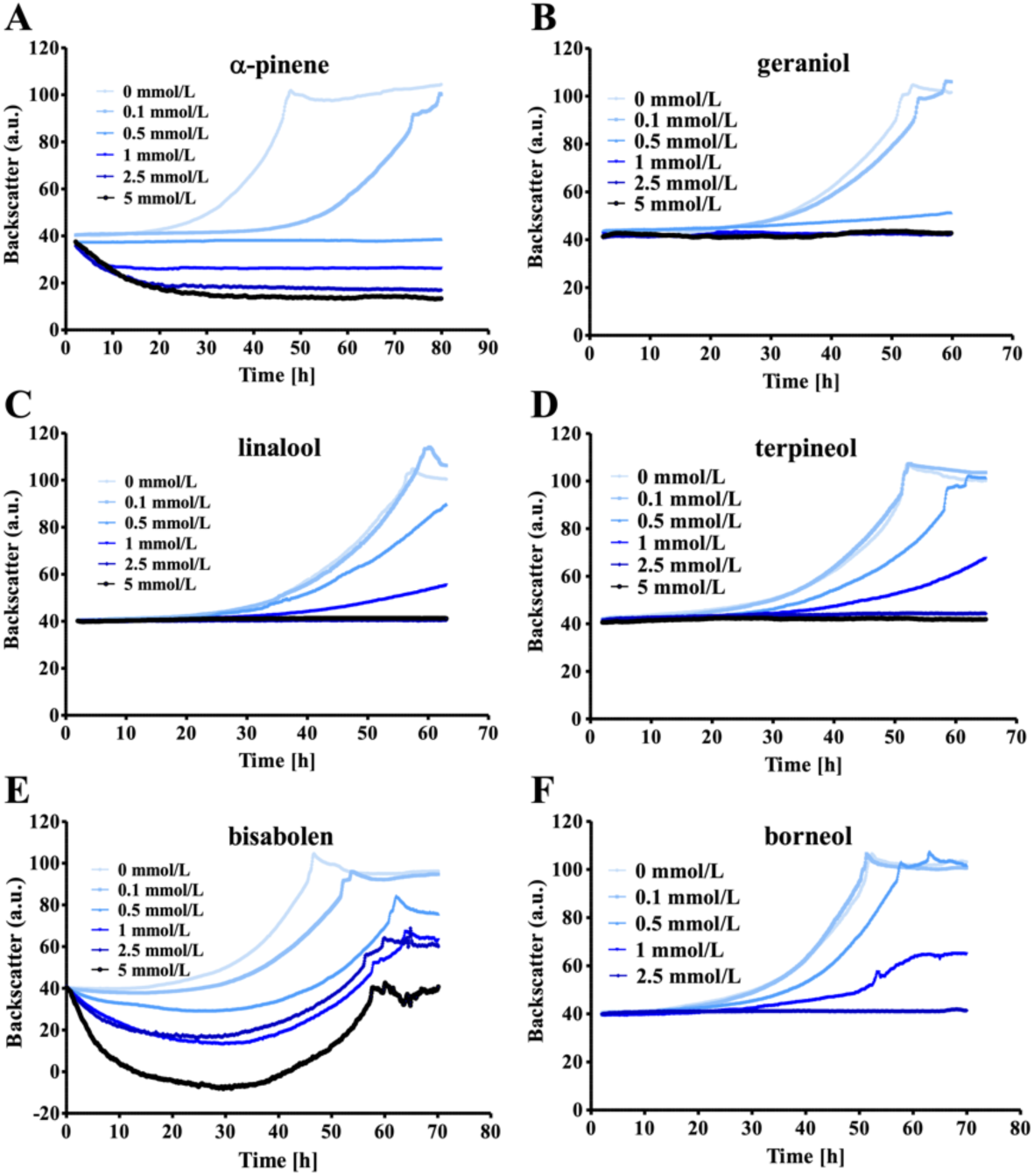
Antimicrobial effect of different isoprenoids on the growth of *M. acetivorans*. Strain WWM73 cultivated on HS medium with methanol and acetate. Linalool (A), terpineol (B), geraniol (C), α-pinene (D), borneol (E) and bisabolene (F) were added in concentrations of 0.1 mmol/L, 0.5 mmol/L, 1 mmol/L or 2.5 mmol/L, respectively. A preculture was inoculated to an OD_578_ of 0.05 into medium containing the different isoprenoids. The cultures were then incubated at 37 °C and shaken at 130 rpm. Cell growth (given in arbitrary units, a. u.) was measured nephelometrically at 515 nm using backscatter measurement in the Cell Quantifier (from AquilaBiolabs). The diagrams indicate mean values and their standard deviations (as error bars) from experiments performed in (at least) triplicates.

Inspired by these results, we decided to introduce a bisabolene synthase into *M. acetivorans* to produce this sesquiterpene as a proof of principle for a more complex isoprenoid. After introduction of an episomal vector containing the gene for α-bisabolene synthase from *Abies grandis* (lacking the N-terminal signal peptide, Table S4) under the control of a tet-inducible promoter into *M. acetivorans*, the resulting strain WWM73/pAgBIS was transferred three times on HS medium with or without tet induction. On methanol plus acetate, a bisabolene concentration of 0.5 ± 0.2 mg/L of culture was obtained after three days (Fig. 3). After three passages on HS medium with tet and CO, a notable increase in bisabolene production, reaching 1.7 ± 0.6 mg/L (Fig. 3), was observed. The difference between the two energy sources might be due to the fact that *M. acetivorans* produces acetyl-CoA only for anabolic purposes on methanol plus acetate, but as a catabolic intermediate when growing on CO (Schöne et al., 2022). Thus, there should be sufficient educt for isoprenoid production via the mevalonate pathway. Our data indicated the suitability of *M. acetivorans* as production host for the isoprenoid bisabolene. Still, the amounts of bisabolene produced were not competitive to known bacterial production systems. Thus, we aimed to optimize the strain and the cultivation procedure.

**Figure 3:**
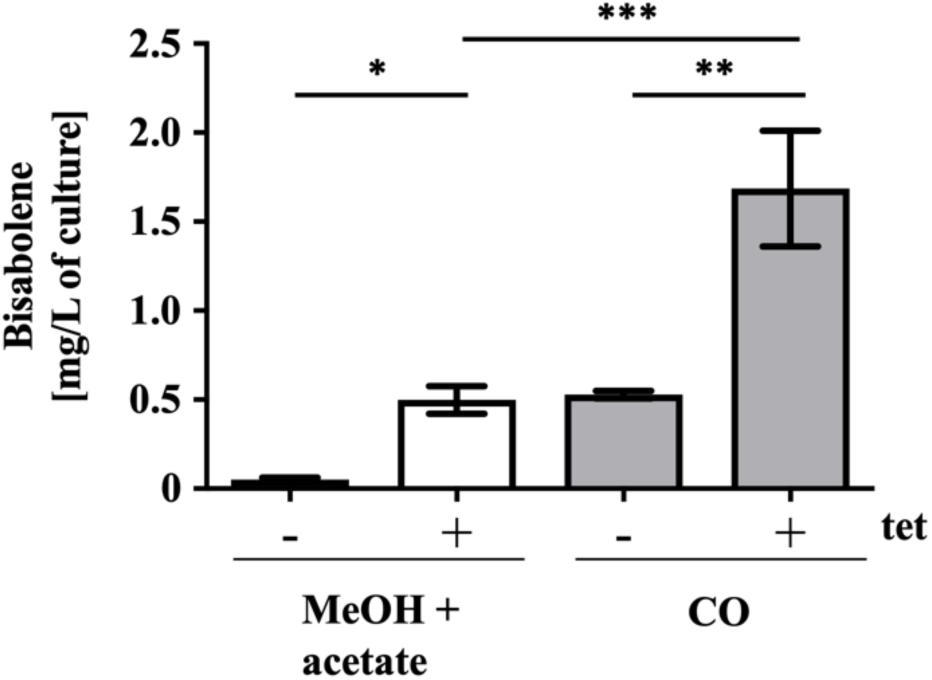
Bisabolene formation in *M. acetivorans* WWM73/pAgBIS. Gas chromatographic analysis of the organic dodecane phase of cultures grown on methanol plus acetate or on CO, for three days in the presence (+) or absence (-) of 100 μmol/L tet. The amount of bisabolene in mg per L culture volume is shown. The bars represent mean values and their standard deviations (as error bars) from experiments performed in (at least) triplicates; comparison was performed using a One-Way ANOVA with Tukey’s post hoc testing. * p≤0.05, ** p≤0.01, *** p≤0.001, ns: not significant.

### Increasing bisabolene production in M. acetivorans

First, we were interested in optimizing induction and growth conditions of WWM73/pAgBIS. After transferring the WWM73/pAgBIS strain twice into tet-containing media, maximum bisabolene yield was achieved, as was previously shown for a PmcrB(tetO1)-*uidA* fusion (Guss et al., 2008). Thus, this procedure was adopted for all subsequent experiments. Next, the question was addressed if growth and bisabolene production are coupled, as many biotechnological systems rely on separating growth phase and production phase (Klamt et al., 2018). To this end, tet-induced WWM73/pAgBIS was grown on methanol plus acetate to stationary phase and further incubated (Fig. 4A). Bisabolene production continued after the culture reached stationary phase and increased even further, when methanol and acetate was fed (Fig. 4A, arrows at day 9 and day 19). Employing this “repetitive closed batch” approach, bisabolene concentration could be increased around sixfold (compared to the beginning of the stationary phase, day 4, Fig. 4A) to 10.6 ± 0.6 mg/L after 23 days, when the experiment was terminated. In this experiment a volume-specific productivity of 0.017 +/- 0.008 mg L^−1^ h^−1^ and a specific production rate of 0.14 +/- 0.05 nmol OD_578_^−1^ h^−1^ was determined (Table 1).

**Figure 4:**
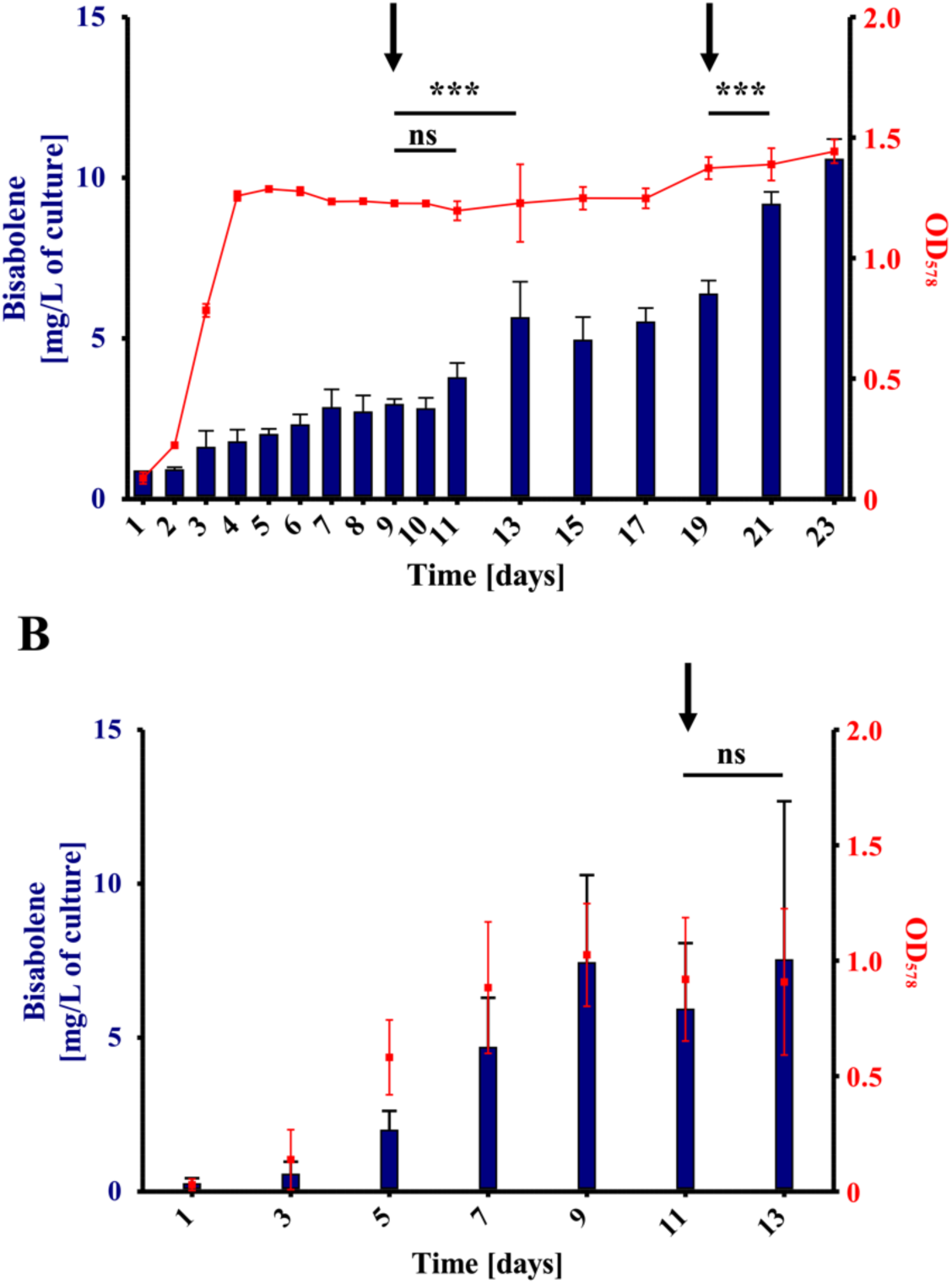
Correlation between growth and bisabolene production of WWM73. Cultures of *M. acetivorans* WWM73/pAgBIS were cultured on methanol plus acetate (A) or on CO (B) in the presence of 100 μmol/L tet. Cultures were overlaid with dodecane (10% v/v) to extract the bisabolene from the medium. Bisabolene in the organic phase was quantified gas chromatographically. The amount of bisabolene in mg per L culture is indicated on the left y-axis (blue bars,). Growth (red squares, right y-axis) was determined photometrically at 578 nm (OD_578_). The arrows indicate the addition of 125 mmol/L methanol and 40 mmol/L acetate (A) or 150 kPa CO (B). Shown are mean values and their standard deviations (as error bars) from experiments performed in triplicates; comparison was performed using a One-Way ANOVA with Tukey’s post hoc testing. * p≤0.05, ** p≤0.01, *** p≤0.001, ns: not significant.

**Table 1:** Reference parameters for bisabolene production of *M. acetivorans* in growth experiments^a^.

| Strain | Energy source | Max. concentration (mg L<sup>-1</sup>) | Volume-specific productivity (mg L<sup>-1</sup> h<sup>-1</sup>) | Production rate (nmol/h) | Specific production rate (nmol OD<sub>578</sub><sup>-1</sup> h<sup>-1</sup>) |
| --- | --- | --- | --- | --- | --- |
| WWM73 | MeOH + acetate | 10.6 +/- 0.5<br>(after 23 days) | 0.017 +/- 0.008 | 0.17 +/- 0.04 | 0.14 +/- 0.05 |
| WWM73 | CO | 7.6 +/- 2.5<br>(after 13 days) | 0.021 +/- 0.012 | 0.62 +/- 0.29 | 0.71 +/- 0.18 |
| MVA231 | CO | 12.2 +/- 1.6<br>(after 11 days) | 0.040 +/- 0.014 | 1.15 +/- 0.23 | 0.97 +/- 0.14 |

a: Data were taken from Fig. 4 (WWM73) and Fig. 6 (MVA231), respectively; rates were calculated between timepoint with linear product increase.

During growth on CO WWM73/pAgBIS produced a maximum concentration of bisabolene of 7.5 ± 2.8 mg/L of culture, which was achieved already after 9 days (Fig. 4B), a concentration reached only after 19 days when methanol plus acetate was the energy source (Fig. 4A). In addition to an increased bisabolene concentration during growth with CO as the sole energy source, the volume-specific productivity was with 0.021 +/- 0.012 mg L^−1^ h^−1^ also slightly higher than during growth on methanol plus acetate. The specific production rate during growth on CO was with 0.71 +/- 0.18 nmol OD_578_^−1^ h^−1^ five times higher than during growth on methanol plus acetate (see Table 1). As addition of CO led to considerable cell lysis, probably due to the acetate and formate produced (Rother and Metcalf 2004), feeding with CO (Fig. 4B, arrow, day 11) had, therefore, no beneficial effect on bisabolene production in *M. acetivorans*. Compared to values for engineered strains with a bisabolene synthase from the literature, the bisabolene concentration obtained in this study from the *M. acetivorans* WWM73 strain cultivated on methanol and acetate was about 20 times higher than that of a non-optimized *Yarrowia lipolytica* strain (carrying a bisabolene synthase gene) cultivated on waste cooking oil as substrate (Zhao et al., 2021). The concentration achieved here is also about twice that of a non-optimized *Synechocystis* strain (carrying a bisabolene synthase gene) cultivated with CO_2_ as carbon source (Rodrigues and Lindberg 2021).

The experiments conducted thus far do not suggest that the cellular level of AgBIS might be limiting the bisabolene yield. Conversely, both the differences in bisabolene yield (Fig. 3) and in rate of production (Fig. 4) suggest that the cellular level of acetyl-CoA can become rate-limiting depending on the growth substrate. The availability of the isoprenoid precursors IPP and DMAPP, i. e., the flux of intermediates through the MVA pathway, seemed another attractive avenue to further increase bisabolene production. We therefore aimed to overproduce enzymes of the MVA pathway in *M. acetivorans*. To minimize unwanted recombination events between the endogenous and the *trans*-genes, those from the closely related *M. mazei* Gö1 were used for this purpose (Boccazzi et al., 2000). The complete MVA pathway comprises nine enzymes, which require a total of ten genes for their production (Table S4). To facilitate cloning, the genes for the pathway were organized into three “modules,” and the coding genes were integrated markerlessly (Pritchett et al., 2004) into one of three established chromosomal integration sites: *ssu* (MVA I), MA3852 (MVA II), and MA2965 (MVA III) (Guss et al., 2005; Sattler et al., 2024). In order to regulate gene expression, the “weakest” tet-inducible promoters, PmcrB(tetO3) or PmcrB(tetO4) (Guss et al., 2008) were placed in front of each construct. The double-integrated strains MVA12, MVA13, and MVA23, as well as the triple-integrated strain MVA231, were generated analogously by re-using the integration constructs after their segregation. All double- or triple-integrated strains containing module III carried a single mutation within gene MM_1763 (D_31_N), which, however, was predicted to not interfere with the activity of the corresponding gene product. Subsequently, the self-replicating plasmid pAgBIS was introduced into these MVA background strains. After the generation of the strains overexpressing enzymes of the MVA pathway, their bisabolene production was determined under tet-induced conditions in cultures grown on methanol plus acetate. None of the strains produced bisabolene to significantly increased concentrations compared to the parent strain (Fig. 5). Only the strain with all enzymes of the MVA pathway from *M. mazei* (MVA231) with 3.2 +/- 0.6 mg/L culture showed increased amount of bisabolene if compared to the WWM73 strain. Thus, overproducing the MVA pathway has only a slight positive effect on bisabolene production under the conditions tested here. To achieve this positive effect, all components of the MVA pathway were required, in contrast to other organisms, where the HMG-CoA reductase, often a rate-limiting enzyme, alone let to increased productivity (Bose et al., 2023; Zhao et al., 2023). The findings support the argument that the availability of IPP and DMAPP is not a critical bottleneck during bisabolene synthesis under the experimental conditions employed. Since acetyl-CoA might already be limiting flux through the MVA pathway with methanol plus acetate as substrate, increasing the amounts of downstream enzymes would not have much of an effect. Another potential reason for this observation could be a limitation in the conversion of IPP and DMAPP to geranyl pyrophosphate (GPP) and/or farnesyl pyrophosphate (FPP), essential intermediates for bisabolene synthesis (Fig. 1). To at least tentatively address this question, the MVA231 strain carrying pAgBIS was cultivated on CO in the presence of tet and the amount of bisabolene determined. At the end of the strain’s growth (day 7) it had produced 8.8 +/- 1.2 mg/L bisabolene (Fig. 6), which is nearly twofold more than the corresponding WWM73/pAgBIS strain under the same condition (compare to Fig. 4 B), which argues for a beneficial effect of the MVA-encoding *trans*-genes. Notably, the MVA231 strain carrying pAgBIS continued to produce bisabolene, to a maximal concentration (without feeding CO again) of 10.16 ±0.02 mg/L of culture. Possibly, in this strain carbon could flow better through the MVA pathway towards bisabolene than towards acetate, leading to delayed cell lysis compared to the WWM73 strain. Additional re-feeding CO resulted in a bisabolene concentration of 12.2 ±2.2 mg/L, the highest concentration achieved (Fig. 6). This concentration is comparable to that achieved on methanol plus acetate (Fig. 4). However, as it took less than half the incubation time (11 versus 23 days, compare Fig. 4A and Fig. 6), increasing the cellular level of acetyl-CoA (via growth on CO) and of MVA pathway enzymes (via expressing *M. mazei* genes), substantially improved bisabolene production in *M. acetivorans*.

**Figure 5:**
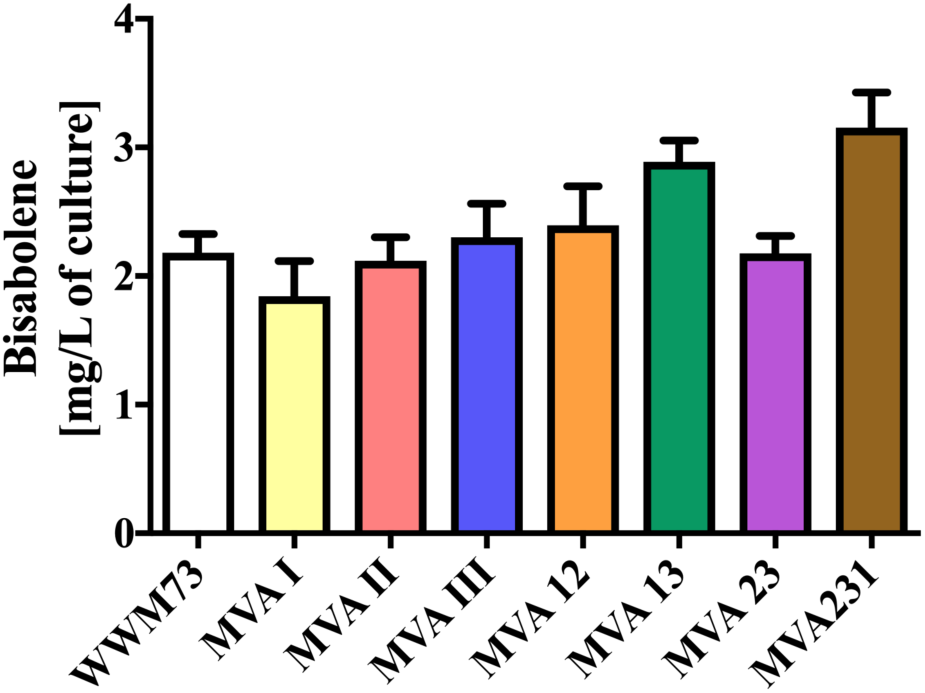
Effect of *M. mazei* MVA pathway enzymes on bisabolene production in *M. acetivorans*. Strains WWM73, MVA I, II, III, 12, 13, 23 and 231 carrying pAgBIS were cultured in HS medium with methanol plus acetate in the presence of 100 μmol/L tet and overlaid with dodecane (10% v/v) for eight days. The amount of bisabolene in mg per L culture is shown. Bars represent mean values and their standard deviations (as error bars) from experiments carried out in triplicates. All mean values were compared against that of the WWM73 strain; comparison was performed using a One-Way ANOVA with Dunnett’s post hoc testing. * p≤0.05.

**Figure 6:**
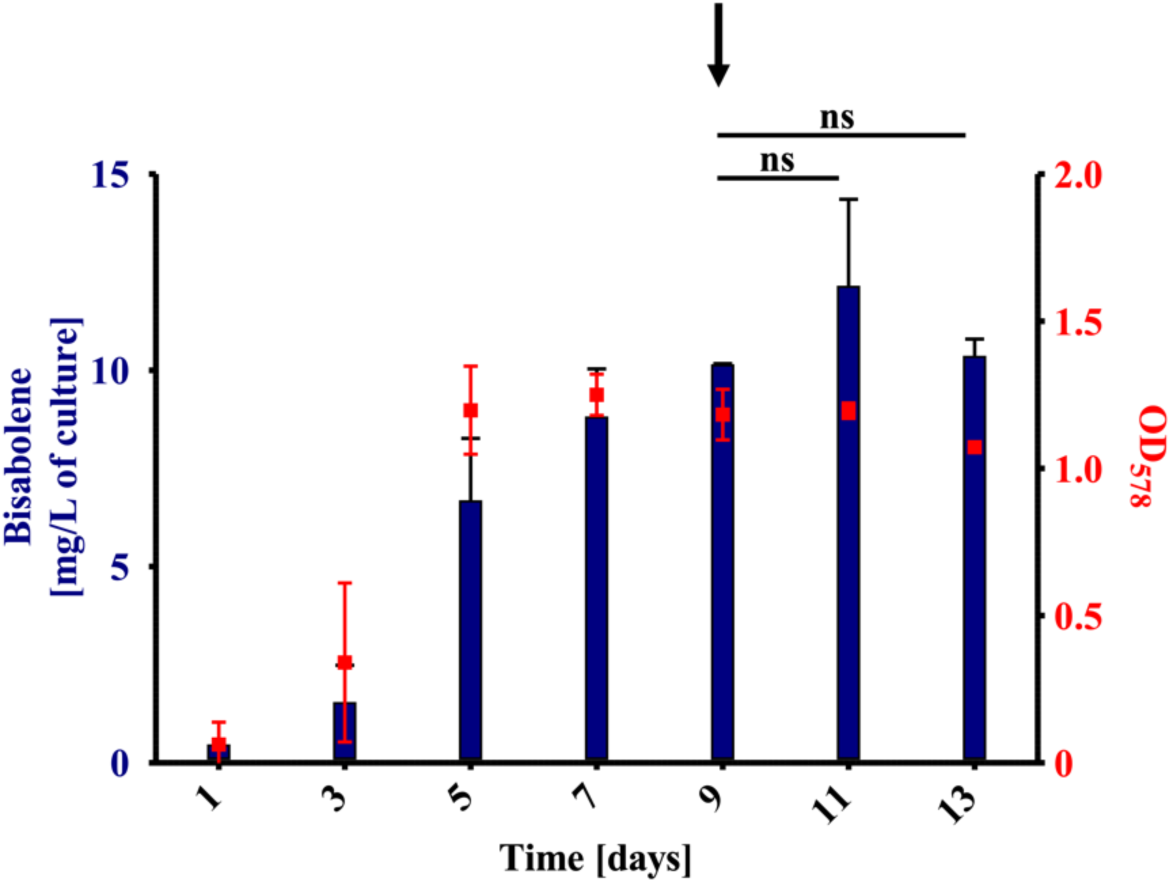
Correlation between growth and bisabolene production of MVA231. Cultures of MVA231/pAgBIS were cultured with CO in the presence 100 μmol/L tet. Cultures were overlaid with dodecane (10% v/v) to extract the bisabolene from the medium. Bisabolene in the organic phase was quantified gas chromatographically. The amount of bisabolene in mg per L culture is indicated on the left y-axis (blue bars,). Growth (red squares, right y-axis) was determined photometrically at 578 nm (OD_578_). The arrow indicates the addition 1.5 atm CO. Shown are mean values and their standard deviations (as error bars) from experiments performed in triplicates; comparison was performed using a One-Way ANOVA with Tukey’s post hoc testing. * p≤0.05, ** p≤0.01, *** p≤0.001, ns: not significant.

In summary, the optimized *M. acetivorans* strain MVA231 on CO produced the highest amount of bisabolene in the shortest time under the experimental culture conditions used, with about 12 mg bisabolene/L culture. *M. acetivorans* cannot yet compete with other production organisms, after extensive optimization, such as *Y. lipolytica* (>280 mg bisabolene/L culture) (Zhao et al., 2021) or *E. coli* (>900 mg bisabolene/L culture) (Peralta-Yahya et al., 2011). To further increase the amount of bisabolene produced in *M. acetivorans*, additional bisabolene synthases from alternative sources will be tested in future studies. Another approach to improve bisabolene production is to introduce terpenoid efflux pumps in order to reduce the cytotoxic effects of bisabolene. This strategy has been successfully applied in *Y. lipolytica* (Zhao et al., 2021; Zhao et al., 2023). Still, the capacity of our genetically modified host to utilize a range of inexpensive substrates, including waste gases (e. g., off-stream gas of the steel industry) instead of complex, and expensive, organic compounds (like fats and sugars), offers the promise of not only higher value added due to lower cost of the feedstock, but also of a step towards a more circular and more sustainable biotechnological application.

## Acknowledgments

This work was supported by a grant from the Bundesministerium für Bildung und Forschung via the MethanoPEP project to M.R. (grant no 031B0851A). The funders had no role in study design, data collection and interpretation, or the decision to submit the work for publication.

## Author contribution

M.R. conceived and supervised the study, A.M. and M.R. designed experiments, A.M., L.V.S., and T.W. acquired the data, all authors analyzed and interpreted the data, A.M. drafted the manuscript, all authors revised the manuscript.

## Supporting Information

**Table S1:**
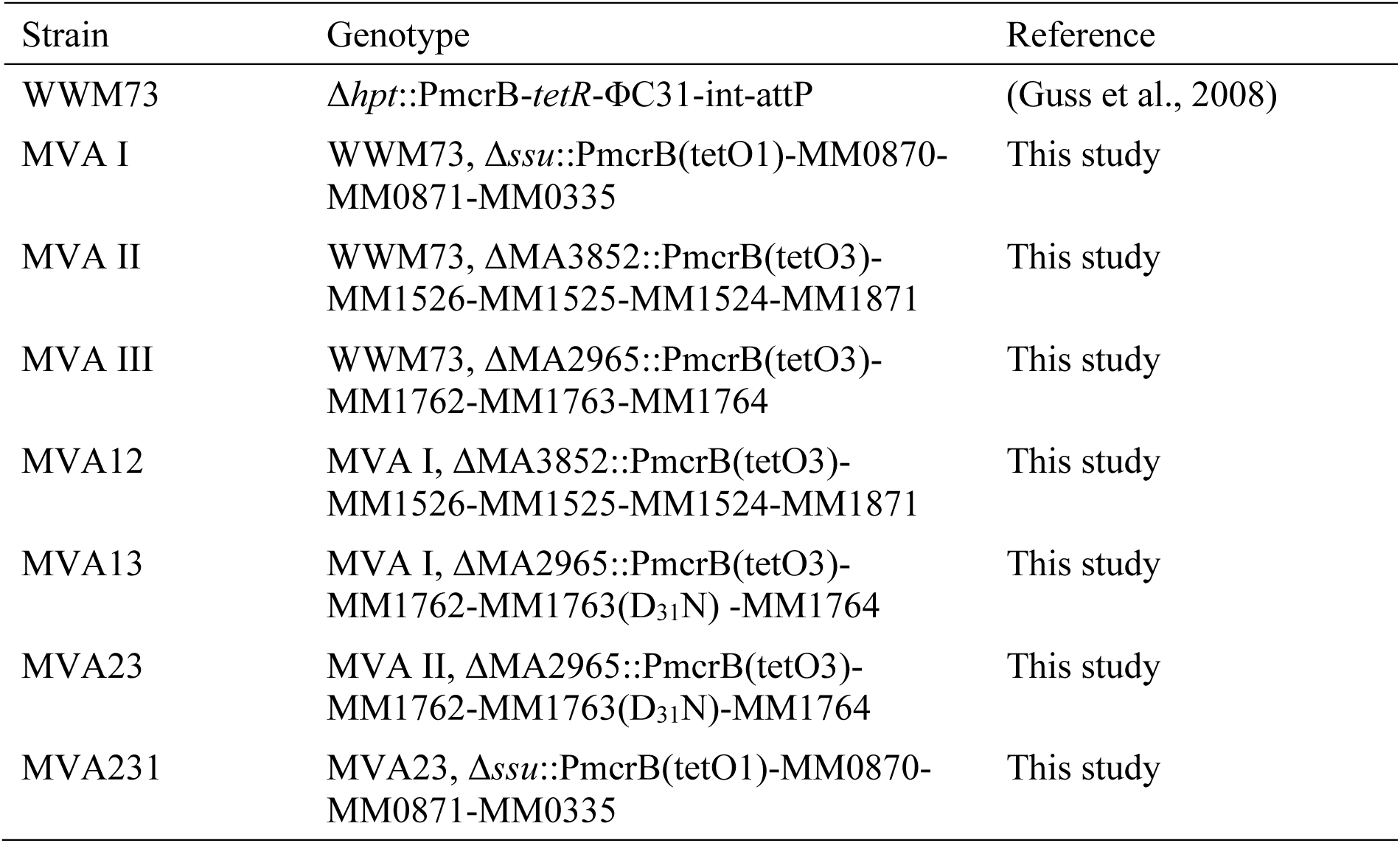
*M. acetivorans* strains used in this study.

**Table S2:** Plasmids used in this study^a^.

| Plasmid | Genotype and/or construction | Reference |
| --- | --- | --- |
| pWM321 | *M. acetivorans*/*E. coli*<br>shuttle vector | (Metcalf et al., 1997) |
| pMR08 | pWM321, $P_{mcrB}\text{-}tetR$ | (Oelgeschläger and Rother 2009) |
| pJK028a | $\phi C31\text{-attB}$ vector (for genomic<br>integration) with $P_{mcrB}(\text{tetO3})$<br>promoter fusion to *uidA* | (Guss et al., 2008) |
| pMssu | Platform vector for chromosomal<br>integration into the *ssu* locus<br>(MA0063-MA0065) | (Sattler et al., 2024) |
| pMA2965 | Platform vector for chromosomal<br>integration into MA2965 | (Sattler et al., 2024) |
| pMA3852 | Platform vector for chromosomal<br>integration into MA3852 | (Sattler et al., 2024) |
| pUC57-BsaI-free | General cloning | Biocat GmbH,<br>Heidelberg,<br>Germany |
| pUC57_tetO1_Pt_APS | Codon optimized α-pinene synthase<br>gene from *Pinus taeda* fused to<br>$P_{mcrB}(\text{tetO1})$ promoter | Biocat |
| pUC57_tetO1_Ms_LMS | Codon optimized limonene synthase gene of *Mentha spicata* fused to PmcB(tetO1) promoter | Biocat |
| pUC57_tetO1_Sc_LIS | Codon optimized linalool synthase gene of *Streptomyces clavuligerus* fused to PmcB(tetO1) promoter | Biocat |
| pUC57/tetO1_AgBIS_His<sub>6</sub> | Codon optimized bisabolene synthase gene (with 3' His<sub>6</sub>-tag encoded) of *Abies grandis* fused to PmcB(tetO1) promoter | Biocat |
| pUC57_tetO4_MVA I | Artificial operon of the *M. mazei* genes MM_0335, MM_0870 and MM_0871 fused to PmcB(tetO4) promoter | Biocat |
| pUC57_tetO3_MVA II | Artificial operon of the *M. mazei* genes MM_1526, MM_1525, MM_1524 and MM_1871 fused to PmcB(tetO3) promoter | Biocat |
| pJK028a_MVA III | MM_1762, MM_1763 and MM_1764 from *M. mazei* (PCR amplified) fused to PmcB(tetO3) | This study |
| pAPS | PmcB(tetO1)-APS fusion (pUC57_tetO1_Pt_APS) moved to pMR08 | This study |
| pLMS | PmcB(tetO1)-LMS fusion (pUC57_tetO1_Ms_LMS ) moved to pMR08 | This study |
| pLIS | PmcB(tetO1)-LIS fusion (pUC57_tetO1_Sc_LIS) moved to pMR08 | This study |
| pAgBIS | PmcB(tetO1)-AgBIS fusion PCR amplified, moved to pMR08 | This study |
| pMssu_MVAI | PmcB(tetO4)_MVA I fusion (pUC57_tetO4_MVA I )moved to pMssu | This study |
| pMA3852_MVA II | PmcB(tetO3)_MVA II fusion pUC57_tetO3_MVA II) moved to pMA3852 | This study |
| pMA2965_MVA III | PmcB(tetO3)_MVA III fusion (pJK028a_MVA III) moved to pMA2965 | This study |
| pMA2965_MVA III D<sub>31</sub>N MM1763 | PmcB(tetO3)_MVA III D<sub>31</sub>N MM1763 fusion (pJK028a_MVA III D<sub>31</sub>N MM1763) moved to pMA2965 (via PCR) | This study |

a: sequences are available upon request.

**Table S3:**
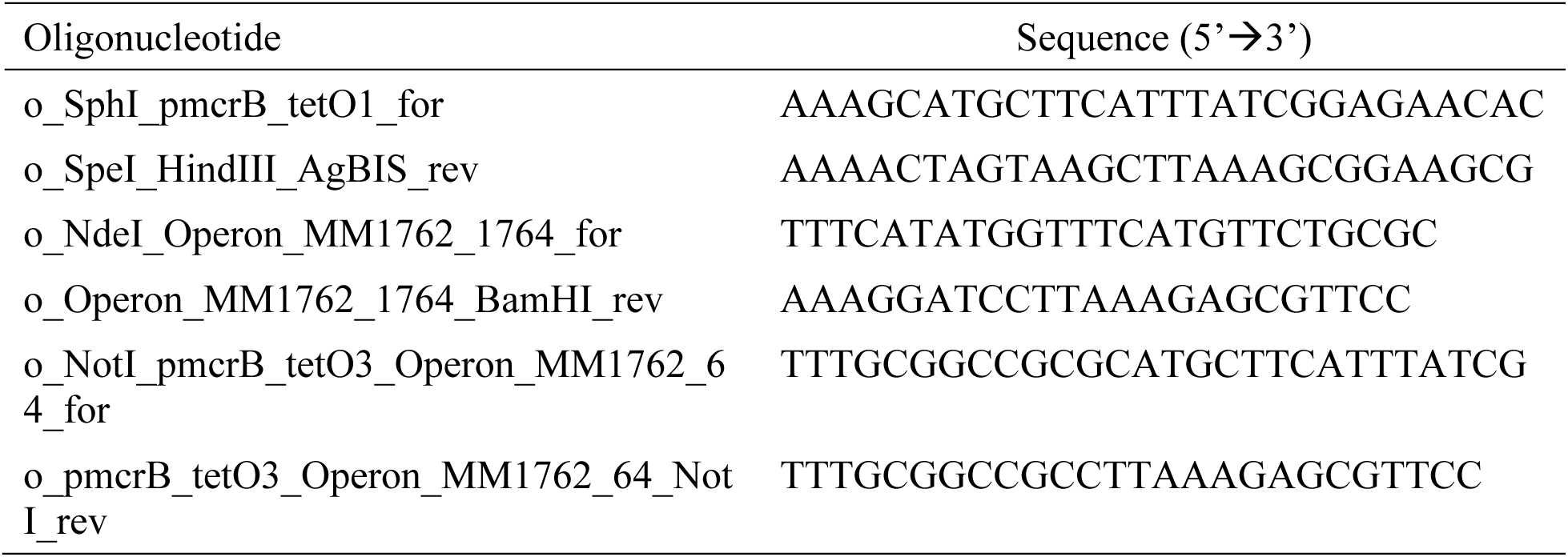
Oligonucleotides used in this study.

**Table S4:** Summary of the *trans* genes used in this study. a: www.ncbi.nlm.nih.gov.

| Name | Encoded enzyme | ORF/accession number<sup>a</sup> | Organism | Comment | Reference |
| --- | --- | --- | --- | --- | --- |
| APS | α-pinene synthase | AF543530 | Pinus taeda | Without plastid transport sequence, codon usage optimized | (Tashiro et al., 2016) |
| LMS | limonene synthase | L13459 | Mentha spicata | Without plastid transport sequence, codon usage optimized | (Wu et al., 2019) |
| LIS | Linalool synthase | CP016560 | Streptomyces clavuligerus | Codon usage optimized | (Nakano et al., 2011) |
| AgBIS | bisabolene synthase | AF006195 | Abies grandis | Without plastid transport sequence, codon usage optimized | (Sebesta and Peebles 2020) |
| MVA I | acetoacetyl-CoA thiolase, HMG-CoA synthase, HMG-CoA reductase | MM_0870, MM_0871, MM_0335 (AE008384) | Methanosarcina mazei | Synthetic operon fused to PmcRB(tetO4) | (Yoshida et al., 2020) |
| MVA II | Pphosphomevalonate dehydratase, anhydromevalonate phosphate decarboxylase, prenylated flavin mononucleotide synthase | MM_1524, MM_1525, MM_1526, MM_1871 (AE008384) | Methanosarcina mazei | Synthetic operon fused to PmcRB(tetO3) | (Yoshida et al., 2020) |
| MVA III | mevalonate kinase, isopentenyl phosphate kinase, IPP:DMAPP isomerase | MM_1762, MM_1763, MM_1764 (AE008384) | Methanosarcina mazei | Operon fused to PmcRB(tetO3) | (Yoshida et al., 2020) |

